# Therapeutic role of Filarial HSP70 in murine models of polymicrobial sepsis and H1N1 Influenza

**DOI:** 10.1101/2021.09.17.460786

**Authors:** Paresh Patel, Shilpa Siddappa, Balachandran Ravindran, Taslimarif Saiyed

## Abstract

Nematodes characteristically modulate effector immune responses by synthesizing and releasing both anti-inflammatory as well as proinflammatory molecules in infected hosts. Pre-clinical studies suggest that immuno-modulatory molecules and synthetic small molecules that mimic parasite products could have therapeutic value to ameliorate tissue damage found in inflammatory diseases. We report here identification of a glycoprotein from filarial parasite, a homologue of mammalian Heat Shock Protein 70 with immunostimulatory attributes. The purified native glycoprotein designated as FHSP70 and its recombinant protein moiety, WFL were found to be TLR2 and TLR4 agonists in vitro in human myeloid cells and induce systemic inflammatory cytokines in vivo. Cecal ligation and puncture (CLP) performed in mice which leads to onset of poly microbial sepsis and mortality could be treated by therapeutic administration of a single dose of FHSP70, along with antibiotics, suggesting its potential as a immunotherapeutic adjuvant for clinical management of Sepsis. Intra-nasal administration of WFL to mice followed by challenge with virulent human Influenza-A virus resulted in decreased viral growth as well as improved survival. The protective effect was demonstrable by both prophylactic as well as therapeutic intranasal administration of WFL. Further, therapeutic administration of WFL by intraperitoneal route 5 days post viral challenge also resulted in significant decrease in viral load in the respiratory tract.

**One sentence Summary:** Systemic administration of a Filarial HSP70 acts as an adjuvant therapy, through immuno-modulation, for improved survival against murine Polymicrobial Sepsis and Viral Infection while its intra nasal administration protects mice prophylactically as well as therapeutically against H1N1 Influenza viral challenge.

## Introduction

Several immunomodulatory molecules have been identified from nematodes *(1, 2)*. These molecules largely have been found to induce anti-inflammatory host responses by generating a dominant Th2 cytokines such as IL-4 and IL-13 *(3–5)* and/or activate macrophages by alternate pathway characterized by decreased generation of inflammatory cytokines viz., TNF-a, IL-1b, IL-6 etc., *(6, 7)*. Nematode mediated suppression of inflammation has been attributed to release of anti-inflammatory molecules by adult parasites and have also been proposed for therapeutic management of auto-immune inflammatory diseases such as IBD, Crohn’s disease and experimental models of arthritis *(8–10)*. Paradoxically, induction of inflammatory host responses are also common features of acute as well as chronic symptoms in systemic helminth infections *(11)*. Growth and development of nematodes in their mammalian hosts have been proposed to depend on induction of inflammatory host responses suggesting presence of inflammatory molecules by worms *(12)* Molecules with inflammatory attributes that activate innate immunity contributing to virulence of parasites in infected hosts have also been reported although less frequently *(2, 13)* Homologues of mammalian alarmin classified as a Danger Associated Molecular Patterns (DAMP) which have been associated with virulence of pathogens have been identified in several eukaryotic parasites (14) and reviewed in (13).

Approaches using molecules such as Eritoran (lipid A derivative) and recombinant anticoagulant molecule, Xigris (Activated Protein C) and other anti-inflammatory molecule have been attempted as Sepsis therapeutic, but, without success in clinical trials *(15, 16)*. Failure of clinical trials for anti-inflammatory molecules that appeared promising in pre-clinical studies has been attributed to induction of immune paralysis in patients with septic shock/severe sepsis rather than hyper-inflammation and cytokine storm observed in Sterile Inflammatory Response Syndrome (SIRS) and early stages of sepsis *(17)* and adverse prognosis is attributable to immune paralysis rather than hyperinflammation. Not surprisingly, several recent studies suggest potential use of pro-inflammatory molecules for Sepsis treatment *(18)*. We had identified a large molecule weight glycoprotein from filarial parasites, designated as AgW displaying high affinity for TLR-4 *(7)*. The glyco moiety was identified to be poly N-Acetyl D Glucosamine (Chitin). We had further demonstrated that its hexa-saccharide, Chitohexose bound to TLR4 and activated macrophages by alternate pathway and blocked endotoxemia *(7)*. In the current manuscript, we describe the protein back-bone of AgW which was found to be a nematode homologue of heat shock protein 70. The molecule designated as FHSP70 was found to be a TLR2 and TLR4 agonist in vitro. In a murine model of polymicrobial sepsis FSHP70 improved survival of animals as an adjunct to antibiotics. More significantly, it also protected mice in a challenge model of viral sepsis. Induction of suppression of innate anti-viral responses in upper respiratory tract by viral pathogens play a critical role in progression of infection leading to lung pathology *(19)*. Intra-nasal application of TLR2/6 agonists that activate the innate immune system have been shown to protect experimental hosts against viral infections *(20, 21)*. In this context the findings being reported here on a single intra-nasal dose of a recombinant filarial HSP70, WFL, inhibiting virulent Influenza-A viral challenge in mice suggests its potential as a candidate anti-viral drug for management of viral influenza in infected humans.

## Results

### Identification of the immunomodulatory glycoproteins fraction of filarial worm

Soluble filarial extracts and their WGA lectin binding glycoproteins (AgW) were found to have immunomodulatory properties *(7)*. However, the active protein(s) that contributed to the immunomodulatory property of AgW were not identified. For this, the AgW protein was purified and subjected to size exclusion chromatography for sub-fractionation. The profile showed multiple peaks, P1, P2, P3 and P4 (Fig 1A). The major protein was found to be eluting in P1. The yields of other peaks P2-P4 were low and they were pooled together and designated as pooled peaks (Pp) for further experimentation. AgW, P1 and Pp activated monocytic cell line, THP-1 and secreted inflammatory cytokines viz., IL6, IL-10, IL-1b, TNF-a and MCP-1 in a dose dependent manner, (Fig 1B) suggesting the inflammatory potential of AgW and purified P1 (but not Pp fraction)

**Fig 1.**
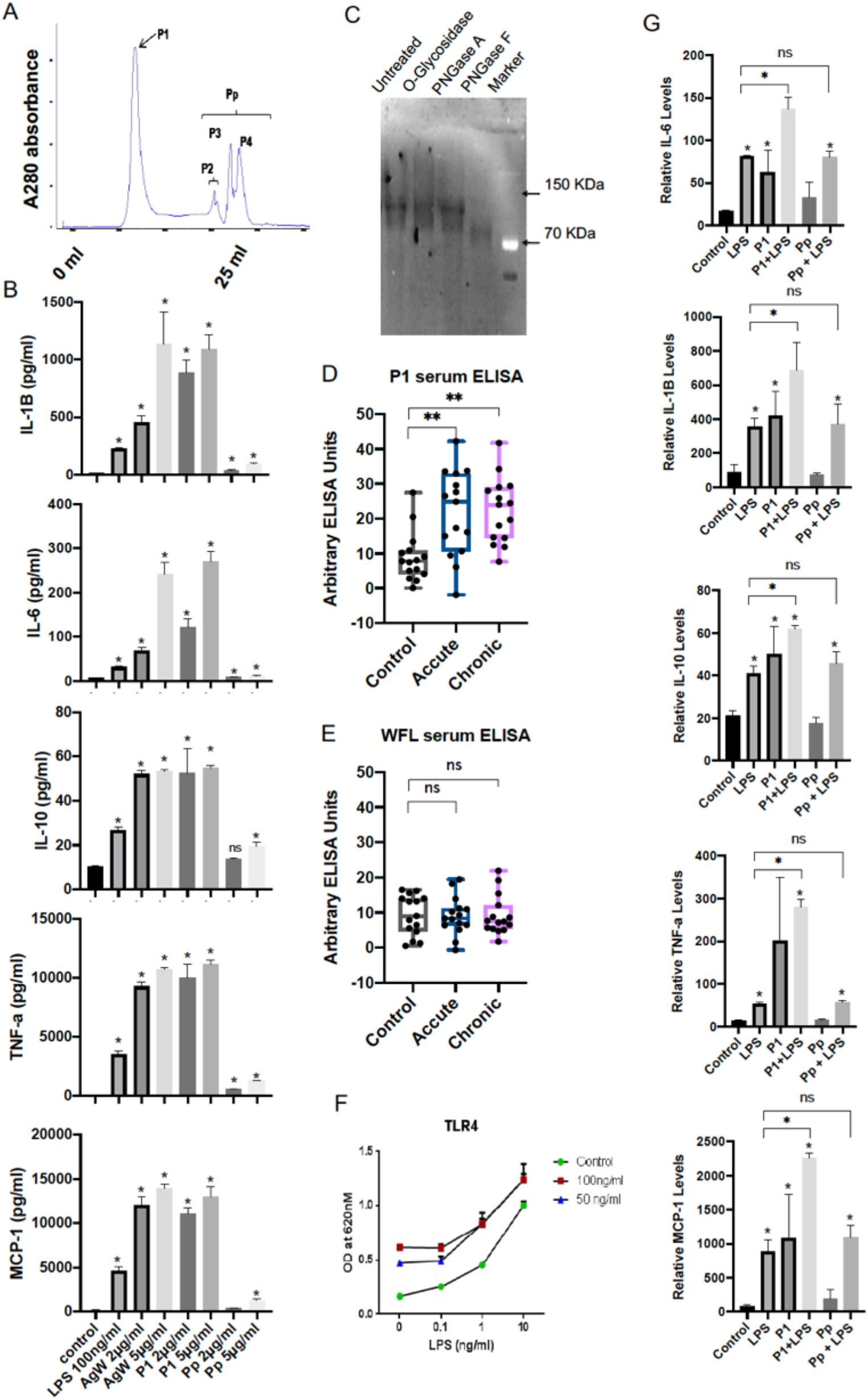
Identification and characterization of immuno-modulatory fraction in WGA binding filarial soluble extracts. A) Different protein peaks (P1-P4) observed in size exclusion chromatography of the AgW. B) Effect of indicated amounts of purified P1 and Pooled peaks (Pp) fraction on secretion of IL-1B, IL-6, IL-10, TNF-a and MCP-1 protein levels in THP-1 culture supernatants. C) Western blot of P1 protein fraction treated with indicated deglycosylating enzymes (biotinylated WGA lectin was used as primary and streptavidin-HRP was used as secondary). B) Alignments and coverage of different peptides generated from proteomic analysis of P1 protein fraction. D) ELISA assay showing the presence of antibodies against native glycosylated P1 or its deglycosylated recombinant form, WFL in serum derived from healthy control, or patients with acute or chronic filarial infection. F) Effect of either 50 or 100 ng/ml P1 protein along with indicated amounts LPS (TLR4 agonist) in HEK-Blue TLR4 reported cells. The NF-kb and AP-1 driven SEAP enzyme activity was assessed by measuring absorbance at 620 nm. G) Effect of purified P1 and Pooled peaks (Pp) fraction on secretion of IL-1B, IL-6, IL-10, TNF-a and MCP-1 protein levels in presence or absence of LPS in THP-1 culture supernatants. *p<0.05, ns= non significant.

### Characterization of P1 protein and its immunomodulatory activity

P1 protein was analysed using western blot with biotinylated lectin before and after treatment with de-glycosylating enzymes viz; O-glycosidase, PNGase A and PNGase F. PNGase F reduced P1 protein to a 70 KDa molecule (Fig 1C) suggesting presence of N-linked glycan. Further analysis by mass spectrometry based proteomic analysis revealed that P1 protein is a homologue of mammalian Heat Shock Protein 70, designated as FHSP70 (Supplementary Figure 1). The gene for FHSP70 was cloned and expressed in a bacterial system to generate its de-glycosylated full Length protein, referred herein after as WFL. Filarial HSP70 sequence is highly conserved among filarial species and displays 77% homology with human HSP70. Given the immune-stimulatory properties of HSP70, we investigated the potential of both Human HSP70 and Filarial HSP70 as a therapeutic molecule against pathogens in this study. Although FHSP70 shares significant homology to its mammalian counterpart, it was essential to investigate its immunogenicity in human filariasis patients with acute or chronic filariasis. Antibodies to native filarial HSP70 (P1) could be detected (Fig 1D) while the reactivity was found to be directed primarily towards carbohydrate moieties since there was no significant reactivity to WFL (Fig 1D) suggesting the protein backbone was either not exposed to the immune system or was not immunogenic owing to its sequence conservation with its mammalian counterpart.

### FHSP70 (P1) activates TLR4 mediated NF-kB and AP-1 activation and synergises with LPS

Filarial glycoproteins have been previously observed to interact with TLR4 *(7)*. To understand whether **FHSP70** activates TLR4 mediated signalling, HEK-Blue TLR4 reporter assay was utilized *(22)* and a dose dependent activation was observed. Activation by **FHSP70** and LPS were synergistic as shown in TLR4 reporter assay and THP-1 cells (Fig 1F and 1G). 50ng/ml or 100ng/ml of P1 was tested in HEK-Blue TLR4 cells along with increasing doses of LPS, where additive activation of TLR4 was observed (Fig 1F). This was further confirmed in THP-1 cells (Fig 1G). The results indicated synergy between FHSP70 and LPS in inducing cytokines and chemokines such as IL-6, IL-10, IL-1b, TNF-a and MCP-1 (Fig 1G).

### Both native FHSP70 (P1) and recombinant FHSP70 (WFL) display comparable TLR2 and TLR4 agonist activity

Immunomodulatory activity of native FHSP70 and WFL were tested using different TLR reporter assays including TLR2, TLR3 and TLR4 (Fig 2A–2C). The results indicated stimulation through TLR 2 (Fig 2A) and TLR 4 (Fig 2C) with no stimulatory activity for TLR3 (Fig 2B). These observations were further confirmed in THP-1 cells using LPS and IL-4 as controls for classical and alternate activation, respectively. P1 and WFL had similar effects on induction of IL-1b, TNF-a, CD274, CCL-13, IL-18 (Fig 2D–2H). Both P1 and WFL significantly increased the levels of CCL-13 (Fig 2F) and CD274 (Fig 2G) and reduced the levels of IL-18 (Fig 2H) consistent with classical activation as seen with LPS.

**Fig 2:**
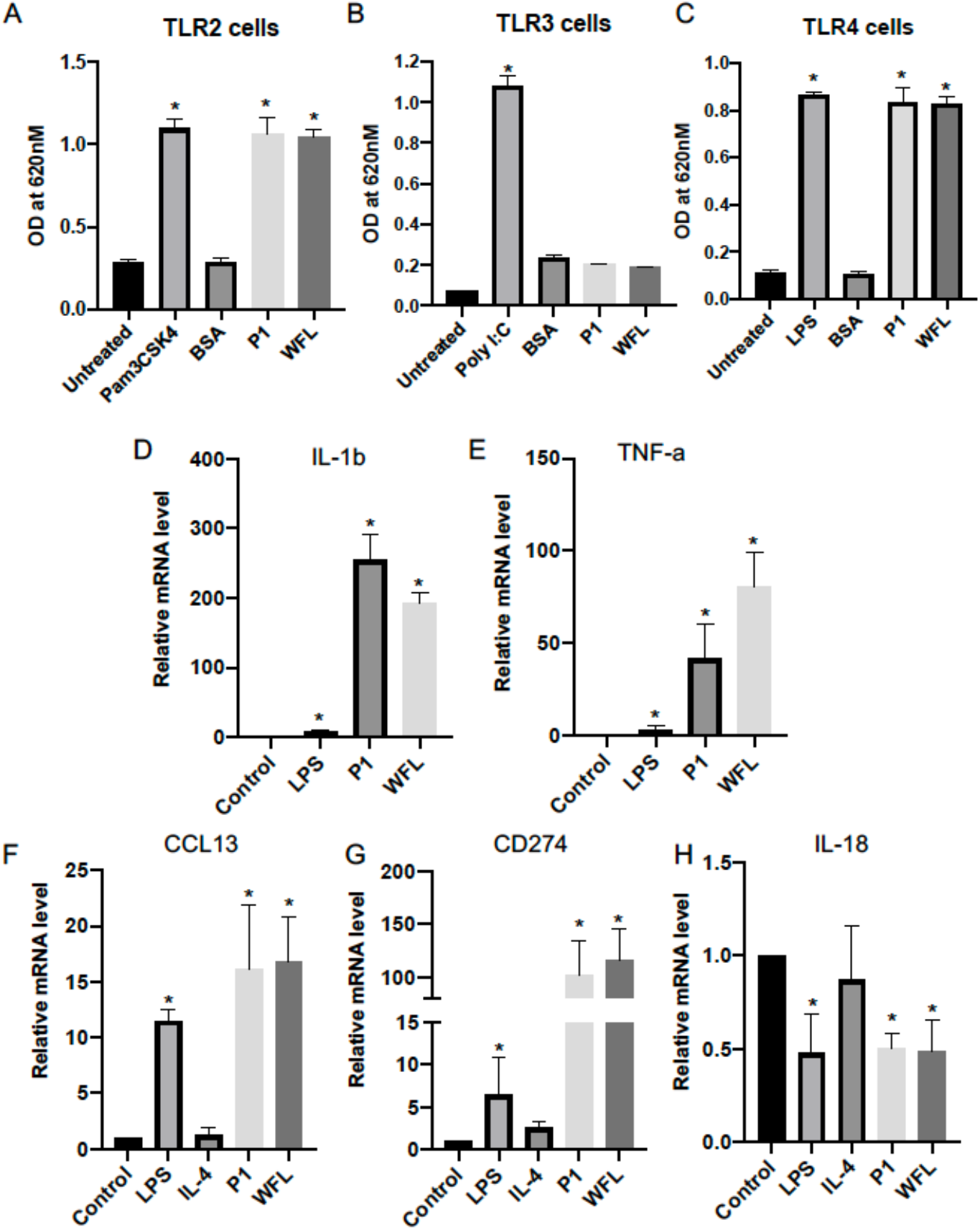
Both native FHSP70 (P1) and recombinant FHSP70 (WFL) display comparable TLR2 and TLR4 agonist activity. Effect of P1 or WFL in A) HEK-Blue TLR2, B) TLR4 and C) TLR3 reporter cells. E-F) Effect of 5ug P1 or 5ug WFL treatment on indicated mRNAs in THP-1 cells 24 hours post stimulation (LPS was used as a control for classical activation and IL-4 was used as a control for alternate activation of monocytes). *p<0.05, ns= non significant.

### FHSP70 (P1) improves survival in polymicrobial sepsis model

Previously, the protective effects of Chitosugars associated with filarial glycoprotein AgW was studied only in a murine model of endotoxemia and not in a polymicrobial sepsis model. In microbial sepsis model, antibiotics are used to control the pathogens while PAMPs continue to activate the immune system leading to uncontrolled immune hyperactivation followed by immune paralysis at a later stage *(23)*. FHSP70 and its recombinant form WFL were tested for early as well as late stage therapeutic effect. A single dose of either 10 ug or 25 ug of test protein was administered intra-peritoneally along with antibiotics at 6 hours post-surgery (Fig 3A) and for late stage intervention, similar doses were administered 24 hours post-surgery and animals were scored for survival (Fig 3B). In the early stage intervention, group treated with 25 ug **FHSP70** showed about 40% survival, while in mice treated with only antibiotics around 20% survived (Fig 3A) and none of the control mice survived post CLP. However, late stage intervention with FHSP70 (P1) protected about 80% of mice in a dose dependent manner (Fig 3B left panel) and Pp treatment showed 50% survival (Fig 3B right panel).

**Fig 3:**
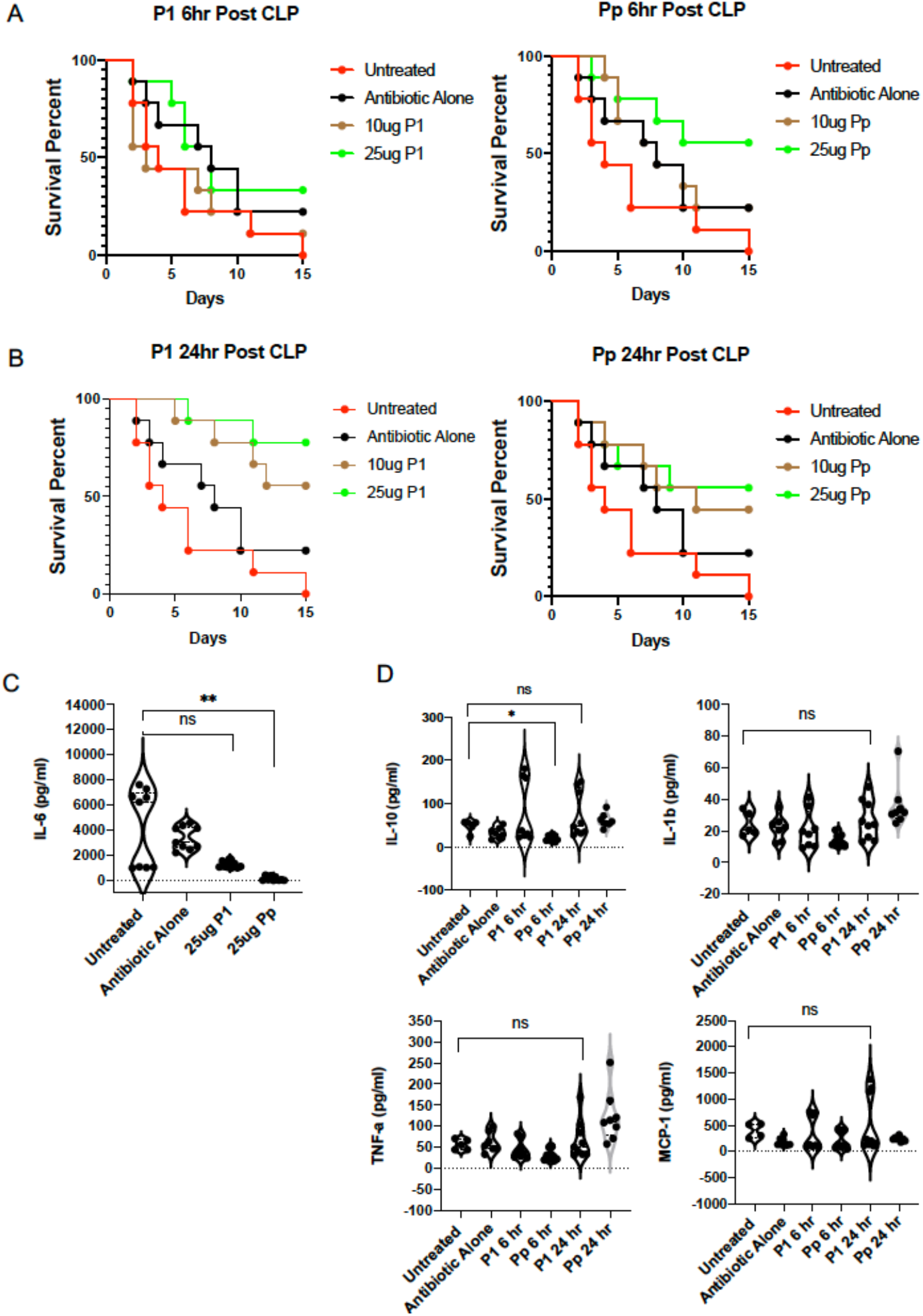
P1 protects mice at late stage intervention, ie; 24hr post CLP and these protective effects are independent pro-inflammatory cytokine regulation. A) Effect of P1 or Pp fractions on survival rate in septic mice when a single dose was administered at 6 hour post CLP along with standard antibiotics. Control ‘untreated’ group did not receive any intervention, while the “Antibiotic alone’ group received a single dose of standard antibiotics at 6 hour post CLP. B) Effect of P1 or Pp fractions on survival rate in septic mice when a single dose was administered 24 hour post CLP. The standard antibiotics were administered at 6 hours post CLP as provided. The same control groups were used as indicated in Fig.3A. C) Effects 25ug P1 and Pp administration on plasma IL-6 levels at 6 hours post intervention (12 hour post CLP) in the groups shown in Fig 3A. D) Effects 25ug P1 and Pp administration on Plasma IL-10, IL-1b, TNF-a and MCP-1 levels 48 hours post CLP in the groups shown in Fig 3A and 3B. *p<0.05, ns= non significant.

Hyperinflammation is a consistent feature of sepsis and cytokines such as TNF-a, IL-6, IL-10, IL-1b and MCP-1 are significantly elevated resulting in a characteristic cytokine storm observed during sepsis. Plasma cytokines were measured to correlate their levels with clinical outcome viz., mortality versus survival in mice undergoing CLP. To understand, whether the protective effects of Pp and P1 are mediated through the control of these cytokine, circulating plasma IL-6 levels were analyzed 6 hours post intervention. Only Pp treated group showed significant reduction as compared to the untreated group (Fig 3C). IL-10, IL-1b, TNF-a and MCP-1 levels were also checked 48 hours post intervention (Fig 3D) and the results showed no significant difference between levels of these cytokines in different groups indicating that the protective effects of late stage intervention with FHSP70 (P1) are independent of regulation of these cytokines.

### WFL is a more potent immunomodulator than its human homolog, HFL

HSP70 is a conserved molecule across species and in-silico docking study revealed that C-terminal domain of filarial HSP70 is important for its interaction with TLR4 (data not shown). The protein sequence of *Setaria digitata*, *W.bancrofti and B.malayi* HSP70 were found to be highly conserved (95-98% sequence similarity), while the human HSP70 displayed only 77% similarity to filarial sequence (Fig 4A). This difference in sequence may contribute to difference in immunomodulatory properties of Human and filarial HSP70. To assess this, a recombinant version of Human full length HSP70, was cloned and purified from bacterial expression system, referred further as HFL. The activity of WFL and HFL were compared in the TLR2 and TLR4 reporter assays. WFL was found to be more potent than HFL in mediating NF-kB and AP-1 signalling (Fig 4B). These findings were consistent when THP-1 cells were used for activation - WFL treatment led to significantly higher induction of IL-6, IL-10, IL-1b and TNF-a (Fig 4C).

**Fig 4:**
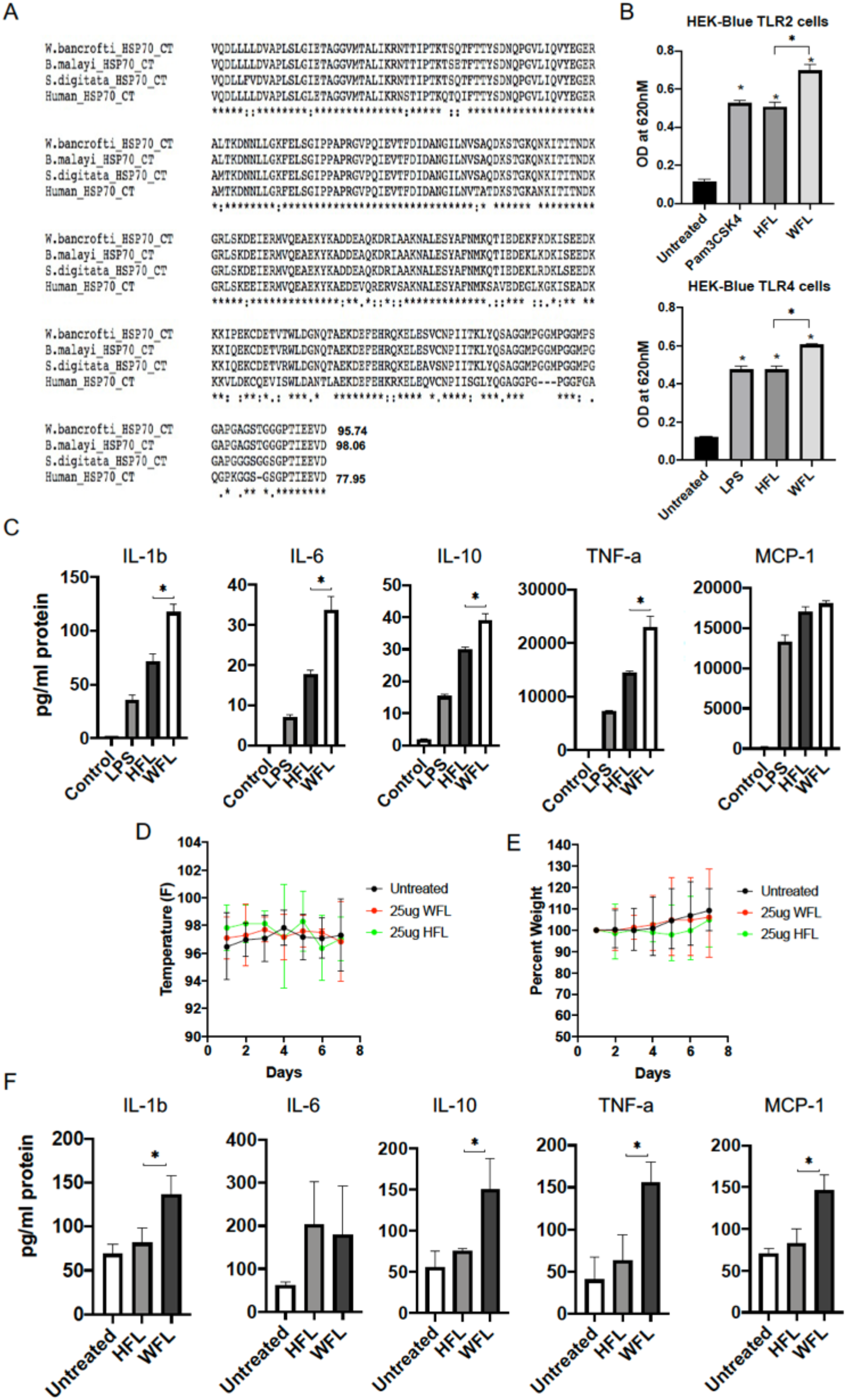
WFL is more potent immunomodulator than its human homolog, HFL. A) The protein alignment of C-terminal (CT) regions of human HSP70 and filarial HSP70s derived from W. bancrofti, B. malayi and S.digitata (The numbers indicate percent similarity when compared to C-terminal region of S.digitata HSP70). B) Effect of equal amounts of WFL or HFL on HEK-Blue TLR2 and TLR4 reporter cells (Pam3CSK4 and LPS were used as positive control in TLR2 and TLR4 assays respectively). C) Effect of WFL and HFL on secretion of indicated cytokines in THP-1. D and E) Effect of single intranasal dose of WFL or HFL on body temperature (D) and weight change (E) in mice. F) Effect of intranasal administration of 25ug of HFL or WFL on plasma levels of indicated innate immune cytokines, 24 hours post application. *p<0.05

### Intranasal delivery of WFL activates systemic innate immune response

The respiratory infection caused by viruses and other pathogens are encountered by the nasal mucosa and innate immune responses by activation of TLRs is proven to be beneficial to the host *(20)*. To investigate if WFL is tolerated in-vivo in mice and is capable of inducing the innate immune system, a single dose of 25ug of WFL or HFL was administered through intranasal route. The body temperature and body weight over 7 days post treatment were measured and the plasma samples were collected 24 hours post treatment. All treated mice were healthy and showed no change in weight and temperature when compared to untreated mice (Fig 4D and 4E). Analysis of plasma samples from these mice revealed the ability of only WFL in inducing a systemic innate immune response when administered by intra-nasal route (Fig 4F) similar to observation made in vitro (Fig 4C), where WFL induced relatively more potent response than HFL.

### WFL treatment reduces viral replication and lung injury in murine influenza model

Lung is the major organ affected during respiratory infections and infections get initiated in the nasal mucosa. Effect of intra-nasal administration of WFL was evaluated in mouse influenza model to monitor viral growth, lung pathology as well as mortality (Fig 5A). In first group, a single prophylactic dose of 25ug was given through intranasal route, 1 day prior to infection. In second group, a single therapeutic dose of 25ug was given, 1 day after the infection to understand the protective effects in early stages of infection. In the third group, single intraperitoneal dose of either 25ug or 50ug was given five days post infection to study the protective effects in viral sepsis. The lungs were harvested fifteen days post infection and visible haemorrhage was observed in mice infected with influenza, which was substantially reduced in the WFL treated mice (Fig 5B). Further histological analysis revealed that immune infiltration and haemorrhage was significantly reduced in all WFL treated group as compared to the infected control mice (Fig 5C). The viral load was measured in the harvested lungs by infectivity assays and the results indicated two to three log reduction in viral titers in treated group when compared to the infected control mice (Fig 5D).

**Fig 5.**
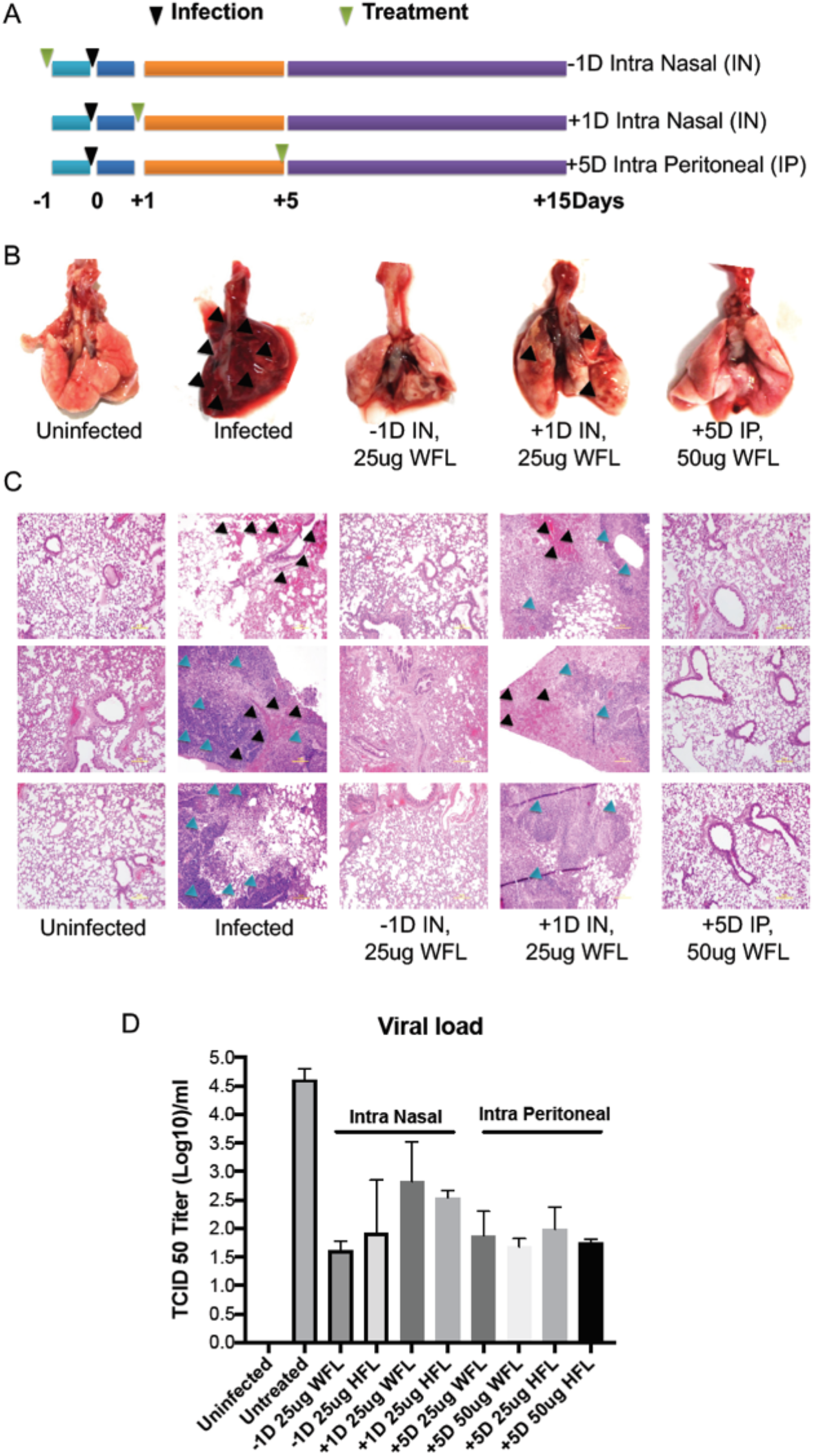
A single prophylactic or therapeutic dose of WFL protects lungs from tissue infiltration and haemorrhage during influenza infection in mice. A) Schematic showing the study plan for influenza infection and different treatment regimes. B) Representative images of the whole lungs at end of study from different groups described in Fig.7A treated with indicated amounts of WFL protein. C) Lung histological images of three independent mice from each group described in Fig,7A, treated with indicated amounts of WFL. D) Influenza viral load in the surviving mice lungs at the end of the study in different groups described in Fig.7A, treated with indicated amounts of WFL of HFL through either intranasal or intraperitoneal route (n=5-9 in each group)

### WFL improves clinical parameter and survival at both early and late stages of influenza infection

The clinical parameter such as temperature, body weight and overall survival were investigated in the different study group described in Fig 5A. The uninfected group showed no change in the temperature over 12 days of the study, while the infected group showed signs of hypothermia with minimum temperature reaching below 94° F on day 6 and the temperature returned to normal in surviving animal by day 10 (Fig 6A–6C). The extent of hypothermia as well as its duration was reduced in the group that received a single prophylactic dose of WFL (Fig 6A). Similar results on hypothermia were observed in the second study group that received a single therapeutic dose 24 hours post infection through the intranasal route (Fig 6B). In the third group, that received an intraperitoneal dose between day 4 and 5 post infection, no significant changes were observed in the temperature between different groups (Fig 6C). Body weight post infection was another clinical parameter that was assessed. Marked reduction in in the body weight was observed in the infected group within 3-4 days post infection as compared to uninfected mice (Fig 6D–6F). The mice that received a single prophylactic dose of WFL or HFL showed no loss in body weight (Fig 6D). Similar results were observed in the group that received a single therapeutic dose 24 hours post infection (Fig 6E). In the late intervention group the weight was reduced for first four days and post intervention on the 5^th^ day with a single dose of 50 ug, both WFL and HFL treated group showed recovery in the lost body weight that returned to baseline levels by day 8 to 9 (Fig 6F). Mice that received the WFL either by intranasal route (Fig 6G and 6H) or by intraperitoneal route (Fig 6I) survived better than control infected mice, WFL being superior to HFL.

**Fig 6.**
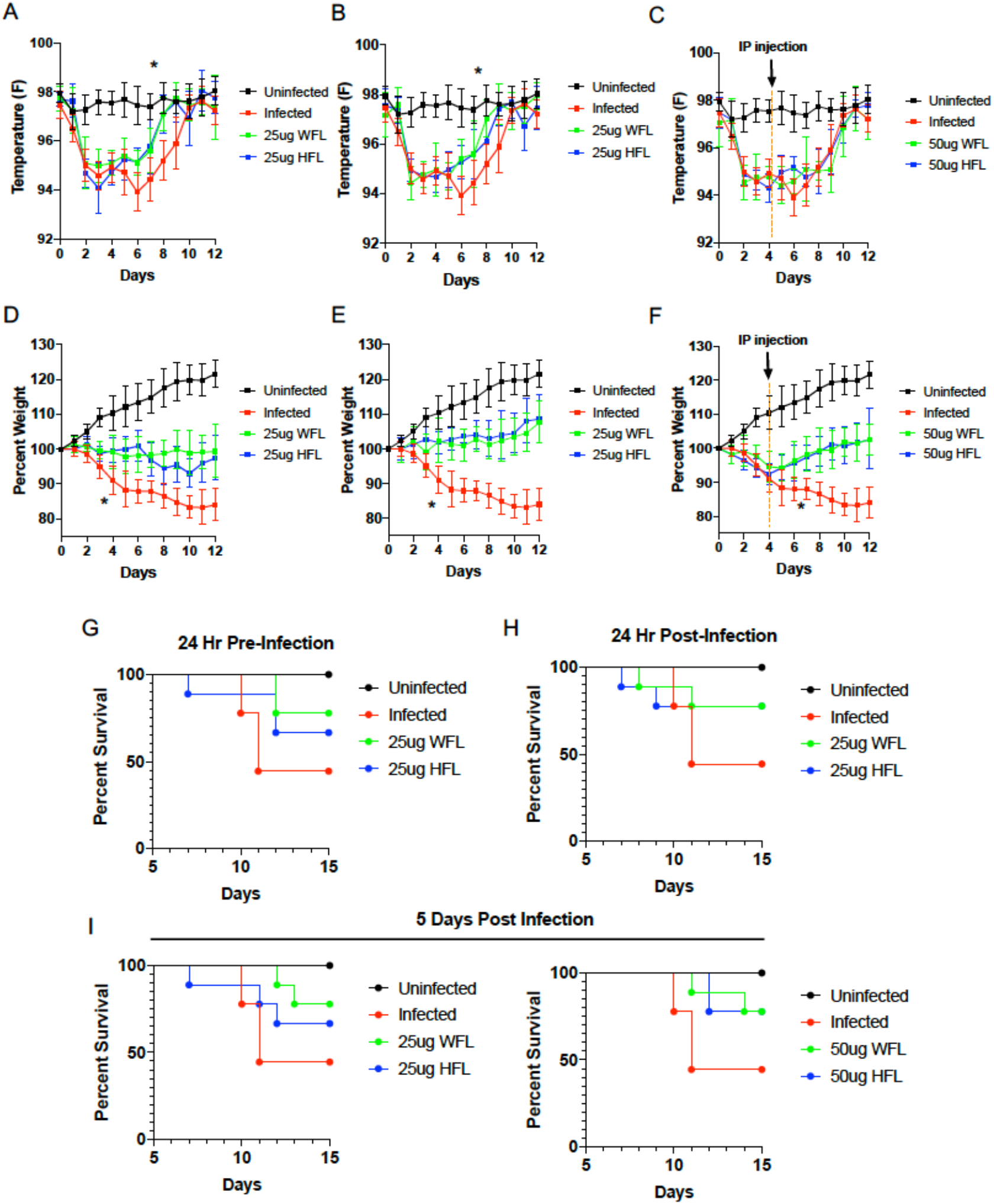
WFL and HFL treatment improves clinical parameters in Influenza infection models. A, B and C) Effect of WFL or HFL on body temperature of mice groups, −1D IN (A), +1D IN (B) and +5D IP (C) as described in Fig. 7A (n=9 in each group). D, E and F) Effect of WFL or HFL on body weight of mice groups, −1D IN (D), +1D IN (E) and +5D IP (F) as described in Fig. 7A (n=9 in each group). G, H and I) Percent survival in mice groups, −1D IN (G), +1D IN (H) and +5D IP (I) described in Fig. 7A (n=9 in each group) that received either 25ug or 50ug of WFL or HFL protein. *p<0.05

## Discussion

The current manuscript reports two striking novel observations on the filarial heat shock protein70, FHSP70 or its recombinant version, WFL. A single intra-peritoneal administration protected mice against mortality from poly microbial sepsis induced by cecal ligation and puncture, a gold standard for evaluating potential drugs in pre-clinical studies for clinical improvement of Sepsis. Second, a single intranasal or intraperitoneal administration of WFL protected mice from developing severe pathology and mortality induced by challenge with influenza-A virus, H1N1 by intra nasal route.

Historically, immuno-modulation using anti-inflammatory molecules as potential drugs have been the mainstay for management of experimental sepsis since induction of unregulated inflammation associated with production of very high levels of inflammatory cytokines has been one of the cardinal features of experimental and human sepsis. However impressive success using anti-inflammatory molecules in pre-clinical studies failed uniformly in subsequent clinical trials *(15, 16, 24)*. This has been attributed to induction of severe immune paralysis observed in patients with septic shock (that follows sepsis) rather than hyper-inflammation per se for adverse prognosis in clinical sepsis *(25)*. This has led to the realisation that use of inflammatory molecules or enhancement of host innate immunity (rather than anti-inflammation) could be more beneficial for management of immunosuppression observed in patients with septic shock. Not surprisingly, inflammasome activation and IL-1b has been found to be beneficial for overcoming immunosuppression in Sepsis *(26)* clinical trials using IFN-g (NCT01649921), GM-CSF (NCT03769844) and IL-7 (NCT03821038) etc are being currently tested *(27–29)* for clinical use.

In the current study, a dominant filarial glycoprotein, a homologues of human heat shock protein70, designated as FHSP70, identified by fractionation and mass spectrometry of previously unexplored glycoprotein (AgW), was found to be a potent immune-modulator and its characterization revealed significant homology of this fraction with HSP70 (Fig 1). It induced significant protection when tested in a murine sepsis model of ceacal ligation and puncture. Our observation that FHSP70 being a pro-inflammatory molecule improves survival against polymicrobial Sepsis only when administered after onset of sepsis (which simulates real life scenario in human patients) and not when administered early (within 6 hours) after CLP procedure, fits into the existing conundrum of pathogenesis of Sepsis viz., immune paralysis observed in later stages of sepsis needing reversal and reprogramming by induction of host inflammatory responses using pro-inflammatory molecules. A recent report on therapeutic benefit of extracellular mammalian HSP72 in a mouse model of peritonitis offers further credence *(30)* since FHSP70 was found to be a TLR2 and TLR4 agonist as shown in this manuscript. Filarial HSP70 homologue was superior to human HSP70 in inducing protection against sepsis possibly attributable to 22% dissimilarity between Human and Filarial HSP70 in their C-terminal sequence. Administration of Mycobacterium w (Mw), another TLR2 agonist, has been reported to be beneficial to patients with severe sepsis patients as shown by improved clinical score and reduced stay in ICU *(31)*.

More significantly, administration of a recombinant filarial HSP70, designated as WFL, a scalable molecule for clinical use by intra nasal route induced systemic inflammation suggesting its translational potential and led us to evaluate its efficacy in an intra nasal challenge model of influenza virus, H1N1 in mice. Although several antiviral drugs have been approved for treatment of Influenza in humans viz., neuraminidase inhibitors, cap-dependent endonuclease inhibitors, humanised single chain antibodies, M2 channel blockers etc., there is an acute need to develop effective therapeutics for management of periodic respiratory viral disease outbreaks in human communities *(31,32)*. In this backdrop, the data presented here on efficacy of WFL is significant - prophylactic as well as therapeutic administration by intra nasal route significantly decreased the viral load, pathology in the upper respiratory tract and mortality (Fig 5 and 6) on challenged with virulent H1N1 strain of Influenza virus. More critically, treatment of infected animals 24hr and even 5 days post viral challenge also mediated measurable protection. These observations offer opportunities for its potential human use as a nasal spray for clinical management of symptomatically infected Influenza patients. Extracellular HSP70 has been reported to innately activate dendritic cells and modulate cytokine production including activation of Interferon regulatory genes (IRGs) and been found to have beneficial effect for cancer treatment *(33, 34)* and the present study is the first to report their potential as an anti-viral molecule with therapeutic efficacy in a pre-clinical model of murine Influenza.

The current study does not however offer significant insights into the mechanism of induction of protection in the nasal tract that generates of functional anti-viral immunity. The observations that filarial HSP70 (free of LPS) innately activate human mononuclear cells in vitro to induce inflammatory cytokines and chemokines systemically when administered by intranasal route suggests its immunomodulatory function. It is however unclear if these host responses were solely responsible for the observed protection in vivo. It is possible that activation of anti-viral innate responses involving type 1 Interferons and pathway of Interferon stimulated genes in the nasal tract could have contributed protection of mice against Influenza-A viral challenge being currently tested in our laboratories. Two reports in literature suggests such a possibility - HSP70 modulates host innate immunity by specifically activating Dendritic cells and macrophages leading to upregulation of several cytokines including anti-viral molecules viz., Type 1 Interferons *(35)*. Plasmacytoid and Myeloid Dendritic cells readily respond with anti-viral host molecules *(36, 37)* on administration of exogenous mammalian *(36–38)*. Reports on Mammalian HSP70 operating through activation of TLR4 and protecting mice against lethal oxidant Lung injury *(39)* and inhibiting influenza viral ribonucleoprotein offer further credence to their potential use as a biotherapeutic agent *(40)*.

Although homologues of mammalian HSP70 have been reported in several eukaryotic parasites *(41, 42)* their immune stimulatory properties and/or their potential as anti-viral drug have not been delineated so far. Immunomodulation by recombinant FHSP70 (WFL) when administered intra-nasally or by intra-peritoneal route mediating functional resistance in mice against clinical sepsis as well as viral challenge suggests its utility for human use. Taken together the data presented here demonstrates therapeutic utility of FHSP70 in pre-clinical murine models for Sepsis, presenting a strong case for undertaking clinical trials as potent adjuvant to standard of care in Sepsis to increase the therapeutic window in polymicrobial and viral sepsis. Pre-clinical studies to evaluate their efficacy as an intranasal formulation for other respiratory viral diseases such as Bird Flu, Swine Flu, SARS-COV2 etc., could lead to development of a broad spectrum anti-viral agent to be used for prophylactic and/or therapeutic purposes.

## Materials and Methods

### Cell lines and serum samples

HEK-Blue TLR2, HEK-Blue TLR3 and HEK-Blue TLR4 reporter cells (Cat. no.# hkb-htlr2, hkb-htlr3 and hkb-htlr4) were purchased from InvivoGen (USA). Human monocytic cells, THP-1 (ATCC-TIB-202) were purchased from ATCC. The cells were cultured in DMEM or RPMI media and were maintained at 37°C incubator with 5% CO2. The culture media was supplemented with 10% Fetal Calf Serum (Gibco) along with standard antibiotics. HEK-Blue selection antibiotics (InvivoGen, Cat. no.# hb-sel, ant-zn and ant-bl) were added as per manufacturer’s instructions. The study on human filariasis was approved by Institutional Human Ethics Committee (IHEC) of Institute of Life Sciences, which operates under the guidance of regulations of Indian Council of Medical Research (ICMR). Written informed consents were obtained from each of the normal control volunteers, filariasis infected persons and/or their legal guardians before collection of blood samples.

### AgW purification and sub-fractionation

AgW, the WGA binding glycoproteins from the adult Setaria digitata worms, was prepared using previously described protocol *(7)*. The purified AgW was subjected to size exclusion chromatography in ÄKTA pure (GE Healthcare Life Sciences) with the Superdex G75 column. The fractions were collected and lyophilized. The proteins were resuspended and passed through EndoTrap^®^ red endotoxin removal kit (LIONEX GmbH, Germany) and stored in endotoxin free water (Cat. No.#95289-1L, Sigma-Aldrich).

### Bioplex based Multiplex cytokine analysis

The Mutiplex immunoassay for Human and Mouse IL-1b, IL-6, IL-10, TNF-a and MCP-1 were purchased from Bio-Rad. For human cytokines, 0.5×10^6/^ THP-1 cells were seeded in 24 well plate a day before and treated with 100ng/ml LPS or indicated amounts of protein for 24 hours. The culture supernatants were directly used for the assay without dilution. For mice cytokines, the plasma samples from mice were collected 6 or 48 hours post treatment and the diluted 1:2 before assay. The assay was performed as per the manufacturers protocol and the beads were read and analyzed in Bio-Plex^®^ MAGPIX^™^ Multiplex Reader (Bio-Rad).

### TLR reporter assay

2×10^4^ HEK-Blue reporter cells were seeded, in each well of a 96 well plate, in HEK-Blue detection media (InvivoGen, Cat. no.# hb-det3) along with the mentioned amounts of test protein. 10ng/ml Pam3CSK4 (InvivoGen, Cat. no.# tlrl-pms.), 100ng/ml Poly I:C (InvivoGen, Cat. no.# tlrl-pic.), or 10ng/ml LPS-EK (InVivoGen, Cat. no.# tlrl-eklps), were used as a positive control for HEK-Blue TLR2, TLR3 and TLR4 cells respectively. The absorbance (@620nm) was measured 16 hours post incubation using TECAN Infinite^®^ 200 PRO multimode reader.

### Ceacal ligation and Puncture study

The Ceacal ligation and puncture (CLP) study with 7-9 week old C57BL/6 males was conducted, at animal house facility at TheraIndx Lifesciences Pvt. Ltd. (Bangalore, India) in accordance with ethical practices laid down in the CPCSEA guidelines for animal care and use *(43)*. The CLP was performed as described before *(44, 45)*. 500ul saline was administered subcutaneously immediately after surgery. Tramadol (20 mg/kg) was injected subcutaneously for post-operative analgesia. Mice received single IP dose at two different dosages of 10ug and 25ug either 6 hours or 24 hours post CLP. Control untreated group was injected with saline solution.

All group except the CLP alone received a single dose of standard antibiotics (amoxicillin and clavulanate) 6 hours post CLP.

### P1 protein identification

P1 protein was analyzed at C-CAMP’s Mass spectrometry-based proteomics facility using Orbitrap Fusion Tribrid Mass Spectrometer (Thermo Fisher Scientific, USA) using optimized protocol for in-gel and in-solution trypsin digested product.

### Western blot for Glycan analysis

The P1 protein was treated with deglycosylating enzymes, PNgase A, PNgase F and O-Glycosidase (New England Biolabs, USA) as per the manufacturers protocol. The deglycosylated protein was desalted and concentrated using Amicon filter (Merck Millipore, USA). The native protein and enzyme treated proteins were separated through SDS-PAGE and the proteins were transferred to the Nitrocellulose membrane using Trans-Blot Turbo Transfer System (Bio Rad). The membrane was blocked using 1x TBST+ 3% Casein followed by overnight incubation at 4°C with biotinylated Lectin from Triticum Vulgaris (Sigma/Merck Cat. No. L5142). The membrane was washed with 1XTBST and further Incubated with Streptavidin PerCP-eFluor710 Conjugated) for 1 h at RT. The images were acquired using iBright Western Blot Imaging Systems (Thermo Fisher Scientific, USA).

### Cloning and purification of WFL and HFL

For Worm full-length (WFL) protein, Setaria Digitata worm was cut into smaller pieces and lysed through TRIzol (Invitrogen, USA) followed by RNA preparation. cDNA was made using RevertAid Reverse Transcriptase (Thermo Fisher Scientific, USA) and the PCR with filarial HSP70 specific primers with appropriate restriction sites (FP: GCAAGATCTATGTCAAAGAACGCAATC and RP: TGAGCGGCCGC CTAATCAACTTCTTCAAT) was performed using Phusion polymerase (Thermo Fisher Scientific, USA). The amplicon was cloned in pET28a vector that adds a His tag at the N-terminal. The human full length (HFL) gene sequence was subcloned from pcDNA5/FRT/TO HIS HSPA1A (a gift from Harm Kampinga, Addgene plasmid # 19537; http://n2t.net/addgene:19537; RRID:Addgene_19537)*(46)* in the pET28a vector. The pET28a-WFL and pET28a-HFL clones were transformed in E. Coli BL21 strain; the cells were induced with IPTG overnight at 18°C followed by sonication and purification using Ni-NTA resin (Cat. No # 786-940, G-Biosciences, USA). The purity was confirmed by SDS-PAGE and the quantity was measured using NanoDrop 2000 (Thermo Fischer Scientific, USA). The pure protein was passed through EndoTrap^®^ red endotoxin removal kit (LIONEX GmbH, Germany) followed by lyophilization.

### P1 and WFL ELISA with Filarial patient samples

2ug/well of purified P1 or WFL protein was coated overnight in a 96 well plate. The plates were washed with wash buffer (1xPBS + 0.2% Tween 20), with shaking (@350 rpm) and blocked with blocking solution (5% Non-fat dry milk + 0.2% Tween 20 in 1xPBS) for 1 hr at RT with shaking. The plates were washed 5 times after blocking and plasma samples from the endemic control, acute and chronic filarial patients were diluted 1:1 in the blocking buffer and added to separate wells followed by incubation at for 1 hr. HRP conjugated Rabbit anti-Human IgG(H+L) (Cat. No. A18903, Invitrogen, USA) as added to each well as the detection antibody. The HRP amount was detected by incubation with HRP substrate and stop solution (1M Phosphoric acid) followed by measurement of OD@450nm in TECAN Infinite^®^ 200 PRO multimode reader. Archived plasma samples collected for an earlier study *(47)* were used for assaying reactivity to WFL and native FHSP70 as described above.

### RNA preparation and RT-PCR analysis

THP-1 cells treated with the test protein and incubated for 24 hours in a 24 well plate. The cells were harvested and the RNA was prepared using RNeasy Mini Kit (Qiagen, Germany) as per the manufacturers protocol. The cDNA was obtained using RevertAid Reverse Transcriptase (Thermo Fisher Scientific, USA). RT-PCR was performed on QuantStudio^™^ 3 and 5 Real-Time PCR System (Thermo Fisher Scientific, USA) using KAPA SYBR^®^Fast qPCR Master Mix (Kapa Biosystems).

### Influenza model study

The Influenza infection study with 6-8 weeks old Balb/c mice was conducted at animal house facility at TheraIndx Lifesciences Pvt. Ltd. under specific pathogen-free conditions in accordance to animal ethics guidelines and approvals laid down by the CPCSEA regulations for animal care and use (44).

The Infection with Influenza A virus (H1N1), strain: A/NWS/33 (ATCC^®^ VR-219^™^) was performed as per previously described protocol *(48)*. Briefly, the 1×10^4^ PFU/mouse in a total volume of 30 μL/mouse in sterile PBS was used for inoculation. The mice were carefully anesthetized using IP Ketamine (60mg/kg) and Xylazine (10mg/kg). The viral inoculum was added through the nostrils and the mice were monitored for any signs of respiratory distress it regained consciousness. The treatment was done in the similar manner, where the test protein at the mentioned amounts was delivered in the nostrils.

### Lung Histological analysis

The lung tissue from the mice were harvested and fixed with 10% buffered formalin and embedded in paraffin. The paraffin blocks were sectioned in 3mm slices and the samples were stained with Haematoxylin Eosin (H&E) stain followed by microscopic analysis for tissue infiltration and alveolar haemorrhage at 100X magnification.

### Viral titers

The work was conducted at TheraIndx Lifesciences Pvt. Ltd. with appropriate safety precautions and approvals. The lungs were harvested and its homogenate was prepared in PBS. MDCK cells were seeded into 96 well tissue culture plate one day prior to the testing so as to get approx. 80% monolayer on the next day. Lung homogenate samples were serially 10-fold diluted in infectious PBS (iPBS) and each dilution was inoculated into cell line (in 96 well tissue culture plate) and incubated for 3 to 7 days at 37°C in 5 % CO2. CPE in the cell culture was observed and virus titer was calculated by Reed-Muench method. Log_10_PFU/g Lung was estimated in each group.

### Bioinformatics and statistical analysis

The protein sequence alignment analysis for the C-terminal domain of filarial worms and the human HSP70 was performed using ClustalW tool (https://www.genome.jp/tools-bin/clustalw). Experimental data were analyzed using GraphPad Prism 8 (GraphPad Software, Inc.). Kruskal-Wallis test and One-Way ANOVA test was used for analysis of cytokines and reporter assay values. Log-rank (Mantel-Cox) test was utilized for survival analysis. P-values, P < 0.05 were considered statistically significant and are denoted by * throughout if calculated P-values were lower than P = 0.05.

## Author contributions

Conceptualization: TS, RB and PP

Methodology: PP and SS

Investigation: PP and SS

Visualization: PP

Funding acquisition: TS, RB and PP

Project administration: TS and RB

Supervision: TS and RB

Writing – original draft: PP

Writing – review & editing: TS, RB and PP

## Data and materials availability

All data are available in the main text and request for the material or the patent details can be made to the

