## Supplementary figures and images for "Therapeutic role of Filarial HSP70 in murine models of polymicrobial sepsis and H1N1 Influenza"

### Supplementary Fig 1

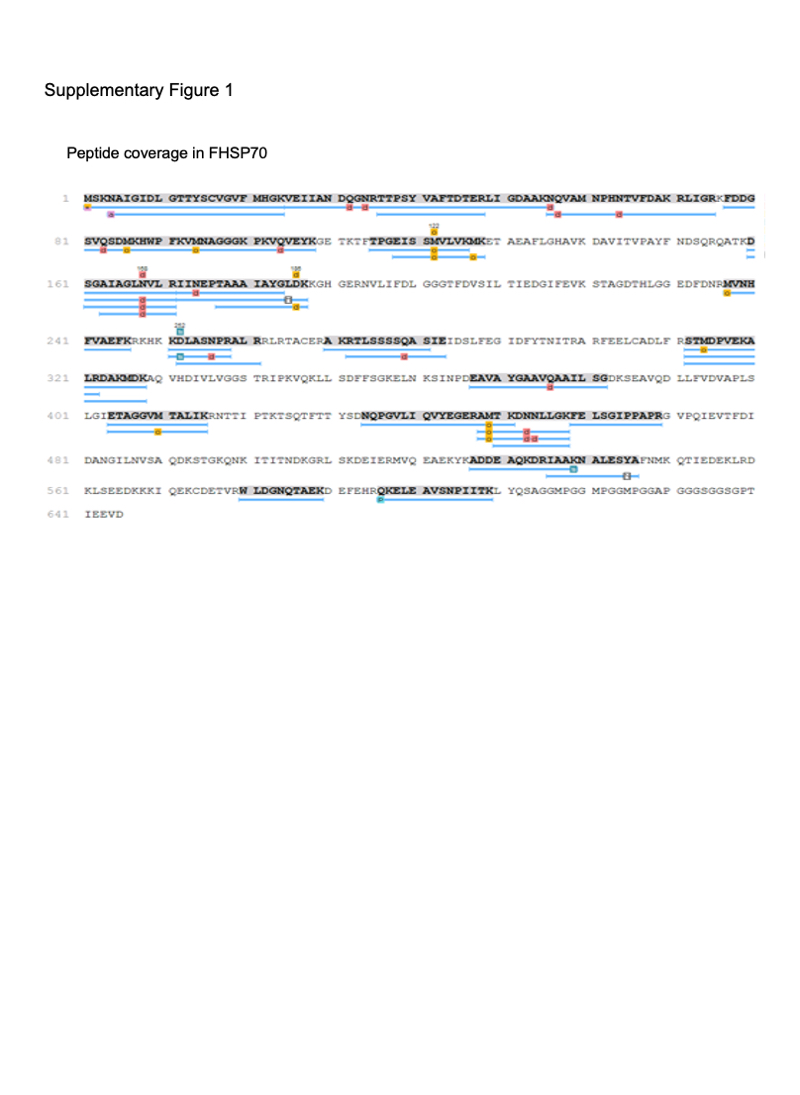
